# The correlated evolution of foraging mode and reproductive in lizards

**DOI:** 10.1101/2022.01.22.477347

**Authors:** Dylan J. Padilla Perez, Dale F. DeNardo, Michael J. Angilletta

## Abstract

Life-history theory suggests that the optimal reproductive effort of an organism is affected by factors such as energy acquisition and predation risk. The observation that some organisms actively search for their prey and others ambush them creates the expectation of different energy needs and predation risk associated with each foraging behavior, the so-called “foraging-mode paradigm”. Although this paradigm has been around for decades, the empirical evidence consists of conflicting results derived from competing models based on different mechanisms. For instance, models within the foraging-mode paradigm suggest that widely-foraging females have evolved low reproductive effort, because a heavy reproductive load decreases their ability to escape from predators. By contrast, a long-standing prediction of evolutionary theory indicates that organisms subject to high extrinsic mortality, should invest more in reproduction. Here, we present the first partial evidence that widely-foraging species have evolved greater reproductive effort than have sit-and-wait species, which we attribute to a larger body size and greater mortality among mobile foragers. According to our findings, we propose a theoretical model that could explain the observed pattern in lizards, suggesting ways for evolutionary ecologists to test mechanistic hypotheses at the intraspecific level.

## Introduction

The foraging behaviors of vertebrates lie along a continuum, ranging from the energetically demanding strategy of searching for prey to the energetically conservative strategy of ambushing prey [2, 3]. During the past 50 years, ecologists have developed a set of hypotheses about how an organism’s foraging mode relates to its life history [4, 5, 6, 7]. These relationships stem from two major assumptions (Figure 1). First, a widely-foraging lizard spends more energy while foraging than a sit-and-wait lizard, but might also consume enough food during an activity season to have more surplus energy [8, 9, 10, 11, 12, 13]. However, a sit-and-wait lizard might have a more diverse diet, because it encounters prey less frequently than a widely-foraging lizard does [14]. Second, a widely-foraging lizard could suffer a greater risk of predation when movements are conspicuous to predators [15]. Risk of predation might also affect the evolution of reproductive effort, because carrying a greater mass of offspring reduces a female’s ability to evade a predator [16]. Both a greater energy supply and a greater mortality risk would select for a genotype that matures at an earlier age and allocates more energy to reproduction, manifested as more or larger offspring [17, 18]. Finally, the load associated with food consumption may compromise the speed of an animal, increasing its vulnerability to predators [19, 20]. In this model, a suite of traits associated with foraging mode would coadapt to the spatiotemporal distributions of prey and predators.

**Figure 1.**
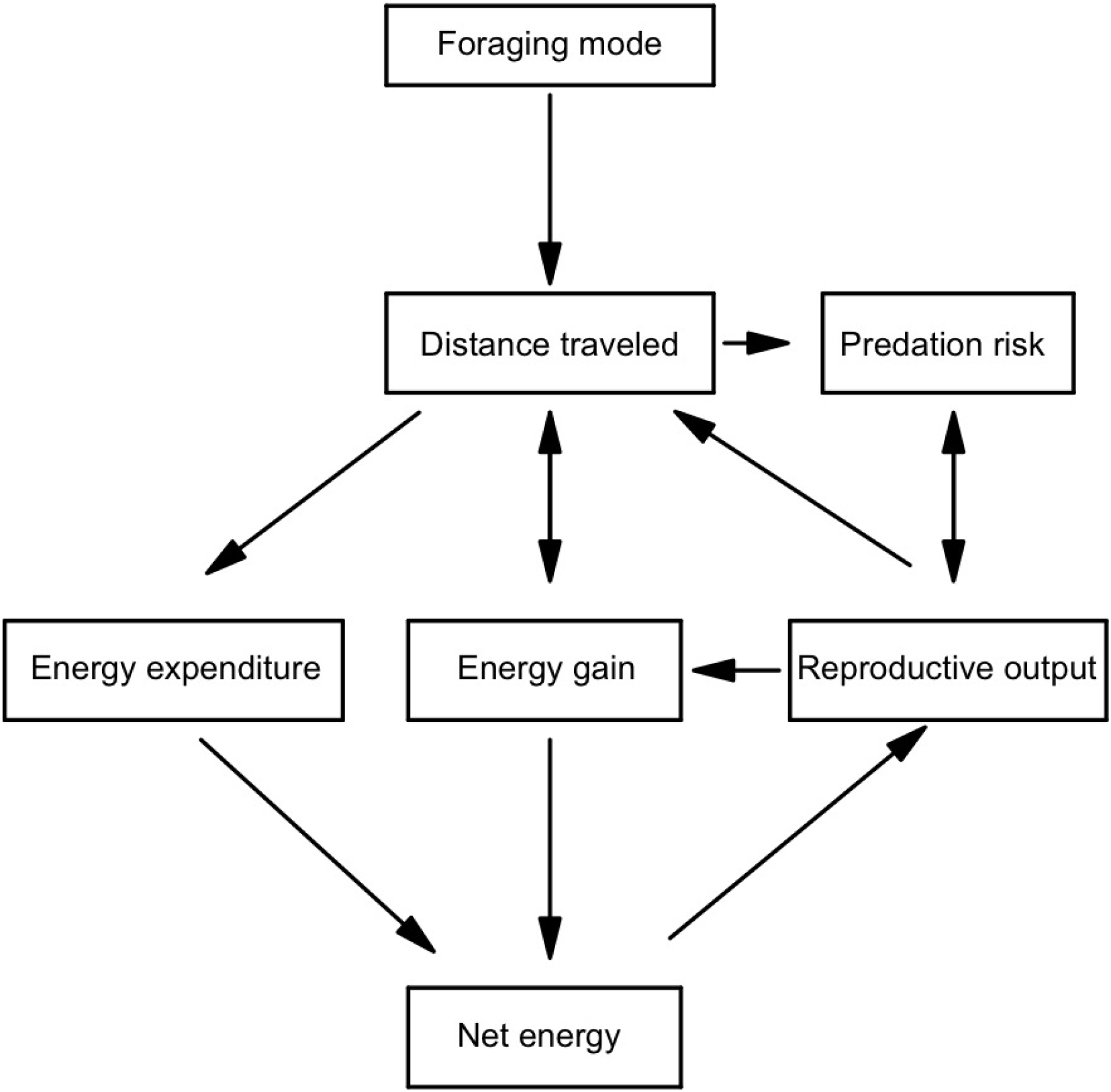
A conceptual model depicting putative relationships among foraging behavior, energetics, predation risk, and reproductive effort. The predicted relationships were derived from theoretical models of life-history evolution (see text for details).

Despite the wealth of conjecture, these hypothetical relationships among foraging mode and other traits have been assessed in only a handful of cases. Ideally, one would isolate each relationship and conduct experiments to quantify the evolution of traits in a controlled environment [21]. However, such data are difficult to gather for many species and will probably remain rare. For instance, limited circumstantial evidence exists for the putative relationship between foraging mode and predation risk. Stomach contents of vipers suggested that widely-foraging species of lizards are more vulnerable to predators than sedentary species [8], but a number of confounding factors can explain this observation as well as foraging mode can. Alternatively, researchers have evaluated the relationship between mortality and life-history traits with no emphasis on foraging behavior. For example, experimental evolution with guppies and fruit flies revealed that genotypes that evolved in risky environments developed more rapidly, matured at a smaller size, and reproduced earlier in life than did genotypes that evolved in safe environments [22, 23]. From these results, we expect that a greater risk of predation among widely-foraging species selects for greater reproductive effort; however, given the scarce evidence, we cannot conclude whether foraging mode generally affects predation risk.

Although experimental data are lacking, comparative methods have been used to explore how foraging mode and life-history traits have evolved. These interspecific analyses have focused mostly on the reproductive of lizards. The earliest analysis of 22 species revealed an interesting pattern: sit-and-wait species had a greater reproductive than widely-foraging species [5]. The authors suggested that widely-foraging species might have to carry fewer or smaller offspring, because moving long distances with a voluminous clutch compromises speed, which in turn decreases the chance of escaping a predator. A subsequent analysis of data for 50 species of lizards supports this result by testing a model in which predation risk increased with increasing reproductive [5]. Roff (2002) extended support for this model in a comparative analysis of 130 species of lizards [24]. However, none of these early analyses controlled for potential phylogenetic correlations that might generate spurious relationships between foraging mode and reproductive [25], especially because foraging mode varies more among families of lizards than within them. A more recent analysis, using phylogenetic comparative methods, failed to detect a significant relationship between foraging mode and reproductive [26]. Thus, the long-standing prediction of a relationship between foraging mode and life-history traits is currently unsupported.

We present the first evidence that the evolution of widely-foraging in lizards was associated with the evolution of greater reproductive effort, specifically among large-bodied species of lizards. This evidence comes from a comparative analysis of foraging mode and reproductive effort comprising 485 species of lizards from 32 families. In this analysis, we inferred the evolutionary history of foraging modes, complementing past reconstructions of ancestral states [27]. In contrast to previous analyses, our study partially supports the prediction of theoretical models of the optimal reproductive effort, paving the way for ecologists to test mechanistic hypotheses at the intraspecific level.

## Materials and Methods

### Sources of data and description of variables

We used published estimates of life history and foraging behavior for 485 species of lizards grouped in 32 families, excluding amphisbaenians and snakes. These data represent a subset of those assembled by ecologists from primary and secondary literatures [28]. We defined reproductive output as the mean product of offspring mass and offspring number. Accordingly, we computed the product of the mean scaled mass index for hatchlings or neonates, 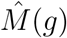, and the mean clutch or litter size for each species of lizards. This product was interpreted in terms of reproductive effort after adjusting for the mass of the parent (i.e., the mean 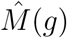 of adult females). The scaled mass index standardizes body mass at a given fixed value of a linear body measurement, based on the scaling relationship between mass and length [29]. We first estimated hatchling mass and maternal mass from values of snout-vent-length (SVL) using published allometric equations [30, 31, 28]. Then, we computed 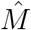 as follows:

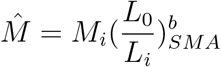

where *M_i_* and *L_i_* are the raw body mass and the linear body measurement of individual i, respectively; *b_SMA_* is the scaling exponent estimated by the standardized major-axis regression of *ln*(*M*) on *ln*(*L*); *L*_0_ is an arbitrary value of *L* (e.g., the arithmetic mean of SVL values for the study species); and 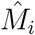 is the predicted body mass for individual *i* when the linear body measure is standardized to *L*_0_. The scaled mass index performs best as a predictor of variation of energy stores, such as fat as well as other body components [29]. Given that we could not find clutch frequencies and ages of first reproduction for all of the widely-foraging and sit-and-wait species in this study, our analysis focused on reproductive effort in a single event rather than the lifetime reproductive effort.

Among species of lizards, widely foraging females and sit-and-wait females have been compared mostly in terms of their relative clutch mass. Relative clutch mass accounts for variation in energy availability among individuals by computing the ratio of reproductive mass to maternal mass, where mass is a proxy for energy. Although relative clutch mass is commonly used as measure of reproductive, regressing this index on female body mass for further analyses might be problematic, because ratios of random numbers regressed against their denominator will yield spurious correlations [32, 33]. By regressing the mean product of offspring mass and offspring number on maternal mass, the slope of the linear relationship can be interpreted as reproductive effort—proportion of mass allocated to reproduction—which enabled us to avoid statistical issues associated with the analysis of ratios.

We defined the foraging mode of a species based on whether it has been reported as ambushing prey (sit-and-wait forager), searching for prey (widely foraging), or using a mixed strategy (mixed). Although this categorization seems somewhat artificial, most species of lizards clearly belong to one of these categories [3, 8]. We focused our analyses only on carnivores, because herbivores do not fit into the classical paradigm of foraging modes [34].

### Statistical analyses

#### Ancestral character state reconstruction and phylogenetic signal estimates

To determine the appropriate model of evolution, we used a set of continuous-time, discrete-state Markov-chain models to sample the character histories from their posterior probability distribution [35], and a time-calibrated phylogeny of squamate reptiles [36]. We rooted the tree with the tuatara (*Sphenodon punctatus*) as the outgroup for our study taxa. Based on previous analyses, we coded this outgroup as a sit-and-wait species [34, 37]. We fitted three different models to our data, using the function *make.simmap* from the “phytools” package of R, version 1.0.1 [38, 39]: 1) an equal-rates model (ER), in which the rate of change between the three states of the character were assumed to be equivalent; 2) an all-rates-different model (ARD), which enables transitions among states to occur at different rates; 3) and a symmetrical model (SYM), which enables pairs of states to change at different rates but changes among all states are theoretically possible. For each model fitted, we estimated the prior distribution on the root node of the tree (*pi* = “*estimated*”) and sampled the transition matrix, *Q*, from its posterior distribution (*Q* = “*mcmc*”). These models sampled the character histories conditioned on the transition matrix (*Q* matrix) and used the phylogenetic tree with annotated tips to create stochastic simulation maps of the potential evolutionary transitions among foraging modes. To describe the variation in reproductive effort among taxa, we plotted bars adjacent to the tips of the phylogeny representing values for each species. We selected the most likely model of evolution based on the Akaike Information Criterion (*AIC*). Lastly, we generated 1,000 trees of the most-likely model and used the *summary* function to count the number of changes in each state, the proportion of time spent in each state, and the posterior probabilities that each internal node was in each state. (see supporting code for detailed information).

We estimated phylogenetic signal (character dispersion on a phylogeny) using Fritz and Purvis’ *D test*, available through the function *phylo.d* in the “caper” package of R, version 1.0.1 [39, 40, 41]. This parameter was calculated for sit-and-wait and widely-foraging species as follows:

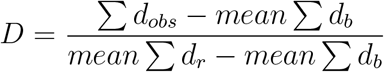

where *d_obs_* equals the number of character-state changes needed to generate the observed distribution of character states at the tips of the phylogeny. The *d_obs_* was then scaled using two null distributions. The first distribution, *d_r_*, comprises d values obtained from permutations, where the number of species with each character state remains constant while the values are shuffled among the tips of the phylogeny. Thus, *d_r_* is the expected distribution of *d* values if character states were randomly distributed among species with respect to phylogeny. The second distribution, *d_b_*, comprises the d values expected if character states were distributed according to a Brownian motion model of evolution. The value of *D* equals 1 if the distribution of the binary trait is random with respect to phylogeny, and exceeds 1 if the distribution of the trait is more overdispersed than the random expectation. The value of *D* equals 0 if the binary trait is distributed as expected under the model of Brownian motion, and is less than 0 if the binary trait is more phylogenetically conserved than expected. The distributions of *d_r_* and *d_b_* were used to assign *p – values* to *d_obs_*. Accordingly, if *d_obs_* is larger than 95% of *d_r_* values, the distribution of the trait would be significantly more overdispersed than the random expectation, if *d_obs_* is less than 95% of *d_b_* values, the character would be significantly more clumped than the Brownian expectation.

#### Effects of maternal mass and foraging mode on reproductive effort

We used Phylogenetic Generalized Least Squares (PGLS) to model the relationship among maternal mass, foraging mode, and reproductive effort. PGLS models enabled us to account for non-independence of the data [25, 42]. To do so, we used the *gls* function from the “nlme” package of R, version 3.1.153 [39, 43]. We included maternal mass and foraging mode as independent variables and the mean product of hatchling mass and clutch size as the dependent variable. Then we fitted multiple models assuming that the species trait values evolved via a Brownian motion model, an Ornstein-Uhlenbeck motion model, and a Pagel’s lambda model using the R package “ape” version 5.6.2. To evaluate the models’ goodness of fit, we used information-theoretic criteria such as *AIC_c_*. We ranked candidate models accordingly and selected the most likely one (lowest value of AICc) for inferences. To control for variation in maternal size among species, we examined models in which maternal mass was standardized 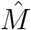. This analysis enabled us to deal with outliers, thus increasing the robustness of our conclusions.

## Results

The evolutionary transitions in foraging mode among species were best described by a model in which rates of evolution were equal among foraging modes, referred to as the equal-rates model (ER). According to the posterior probabilities from stochastic mapping, sit-and-wait foraging is the most likely ancestral state of all lacertilians (Figure 2). Sit-and-wait foraging evolved in two major clades of lizards; near the root of the tree, we found strong evidence suggesting that the ancestor of the basal gekkotans was likely a sit-and-wait predator. Likewise, the ancestor of the more derived iguanians was likely a sit-and-wait predator. During the Jurassic and middle Cretaceous, major transitions from sit-and-wait foraging to widely foraging seemed to have occurred in the ancestors of Scincoidea, Lacertoidea, and Anguimorpha. However, a major reverse transition from widely foraging to sit-and-wait foraging has also occurred during the Jurassic, in the ancestor of iguanians. The *D* test for phylogenetic signal indicated that foraging mode is phylogenetically conserved among lizards (*D* = −0.14, *p*[*D* < 1] = 0, *p*[*D* > 0] = 0.80). Accordingly, closely related species tend to exhibit the same foraging mode, but this similarity decreases as phylogenetic distant increases.

**Figure 2.**
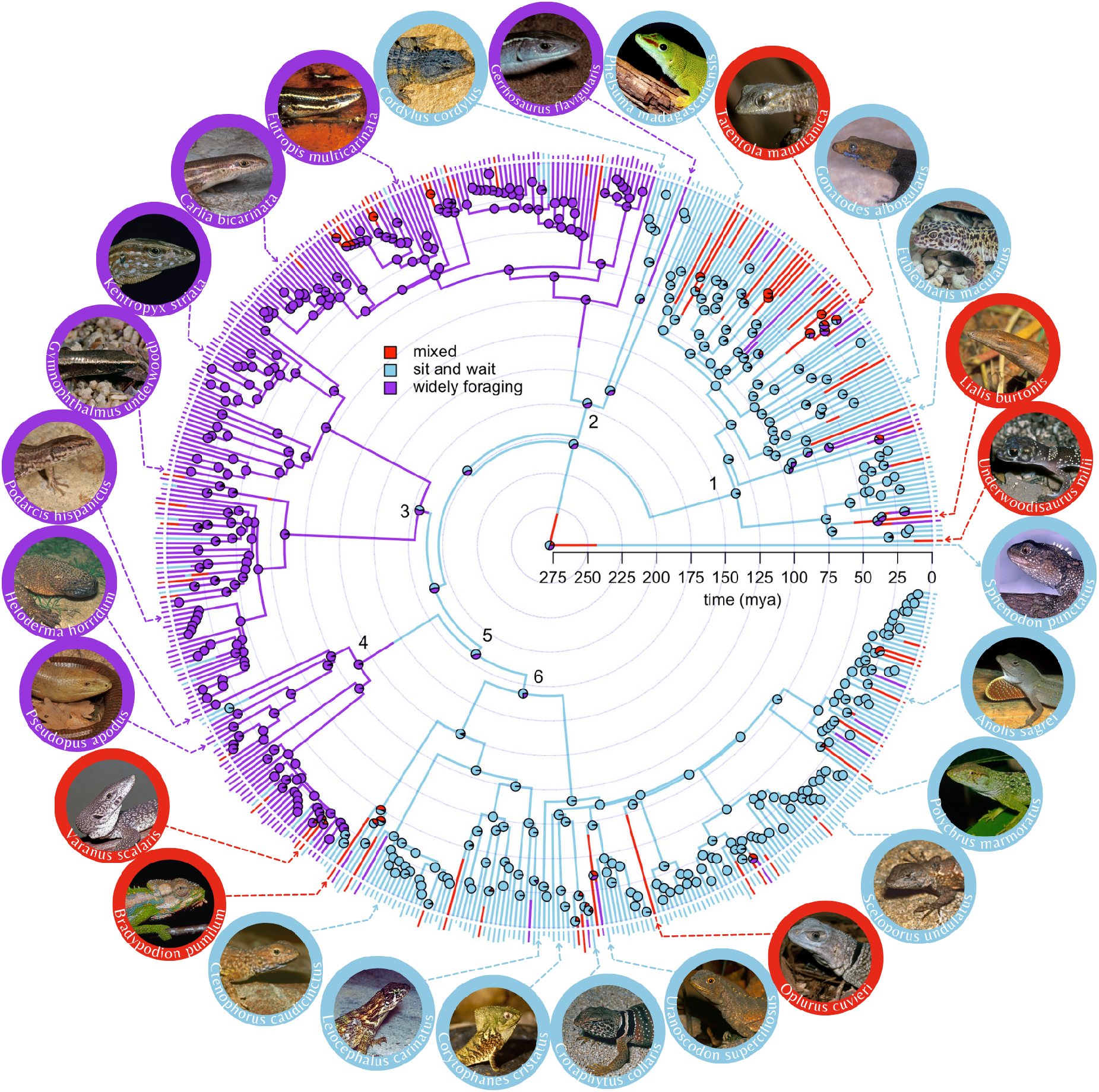
Random sample of stochastic character maps depicting the evolution of foraging mode in 485 species of lizards. Bars at the tips of the phylogeny represent log-transformed values of reproductive effort for all lizards, but not the outgroup, *Sphenodon punctatus*. Pie charts on internal nodes represent posterior probability estimates resulting from 1,000 simulations of the character histories. Major clades are enumerated as follows: 1) Gekkota, 2) Scincoidea, 3) Lacertoidea, 4) Anguimorpha, 5) Toxicofera, and 6) Iguania. Lizard photos by Mark O’Shea.

The evolution of reproductive effort in lizards was driven by an interaction between maternal mass and foraging mode (Table 1). The information-theoretic approach revealed that a model including this interaction was strongly supported (Table 2; Table S1). Widely foraging was associated with the evolution of greater reproductive effort than was sit-and-wait foraging, specifically in large-bodied species of lizards (Figure 3a). Yet, this difference seemed small when hatchling mass and maternal mass were standardized (Figure 3b; Table S2). A model in which mass was standardized, 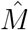, suggested that species adopting a mixed-foraging strategy evolved the greatest reproductive effort (Figure 3b).

**Figure 3.**
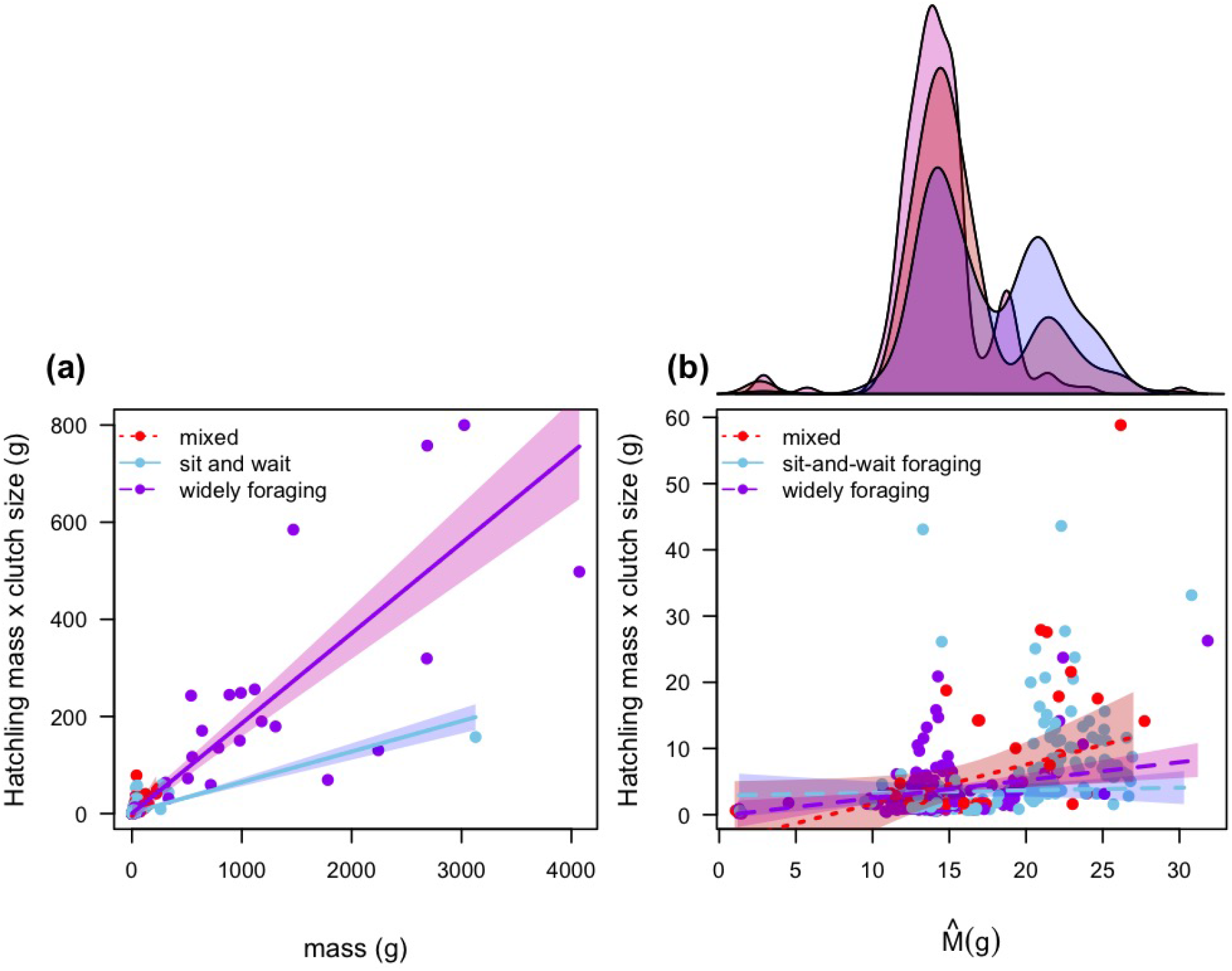
Effects of maternal mass and foraging mode on the evolution of reproductive effort of lizards, as determined by phylogenetic generalized least squares analysis. (a) Difference in reproductive effort among foraging modes assuming body mass as a proxy for energy. (b) Difference in reproductive effort among foraging modes assuming the scaled mass index as a proxy for energy.

**Table 1.** Analysis of Deviance Table (Type III tests) for the most likely model, based on the ranking of *AIC_c_* for potential candidate models.

|  | Df | Chisq | Pr(>Chisq) |
| --- | --- | --- | --- |
| (Intercept) | 1 | 0.005 | 0.941 |
| $\hat{M}_{\_}\text{females}$ | 1 | 4.760 | 0.029 |
| foraging.mode | 2 | 10.372 | 0.005 |
| $\hat{M}_{\_}\text{females:foraging.mode}$ | 2 | 18.104 | 0.000 |

*A colon mark (:) in a model represents an interaction term.*

**Table 2.** Contrast of parameters estimated by the most likely model according to the information-theoretic criteria of selection.

|  | Value | Std.Error | t-value | p-value |
| --- | --- | --- | --- | --- |
| Intercept of widely foraging | 0.222 | 0.384 | 0.578 | 0.563 |
| Intercept of sit-and-wait | 1.804 | 0.186 | 9.698 | 0.000 |
| Intercept of mixed foraging | -5.679 | 3.738 | -1.519 | 0.129 |
| slope of widely foraging | 0.186 | 0.003 | 71.034 | 0.000 |
| slope of sit-and-wait | -0.123 | 0.012 | -10.660 | 0.000 |
| slope of mixed foraging | 0.106 | 0.112 | 0.946 | 0.345 |

## Discussion

Our analysis provides evidence that species of widely-foraging lizards have evolved greater reproductive effort, but only in heavier species (see Figure 3a). Presumably, this pattern stems from the ability of widely-foraging species to harvest and assimilate more resources than sit-and-wait species can. In the field, a widely-foraging lizard spends 32% more daily energy than a sit-and-wait lizard, but this extra energy expenditure is probably paid off with greater daily food consumption [44]. Food consumption may play an important role in determining the reproductive effort of lizards in three ways: 1) promoting follicular growth during the reproductive season; 2) increasing energy stores to initiate reproduction, and 3) reducing age at first reproduction. For example, female vipers that had good body condition early in vitellogenesis produced large litters [17]. Similarly, vipers that gained more mass during follicular growth produced larger offspring. Early reproduction gives offspring sufficient time to mature in the same year that they hatched, which enables them to participate as adults in the subsequent breeding season [45]. Therefore, a widely-foraging species might gain more resources before and during each reproductive event than a sit-and-wait species would, potentially boosting their reproductive effort.

In many species of animals and plants, the reproductive effort increases with increasing body size [46, 47, 48, 49, 50, 51]. This observation indicates that increased size at maturity might evolve according to a reproductive benefit of larger size, balanced by the risk of mortality associated with delayed maturation. Interestingly, the effect of maternal size on the reproductive effort of lizards depended on foraging mode. A model of the evolution of optimal reproduction predicts this pattern [52]. Assuming same daily and seasonal activity durations among species, this model suggests that the total energy accumulated for reproduction (*m*) depends on the time spent foraging (*t*) and maternal size. Thus, the ratio of the energy accumulated for reproduction to the time spent foraging 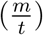 corresponds to the foraging efficiency of an organism. If two organisms have the same foraging efficiency, the smaller one would reach its maximal capacity to accumulate resources at a lower value of *m*, producing fewer or smaller offspring (Figure 4a). However, if a widely-foraging species has a smaller body size but a higher foraging efficiency, it might invest more energy in reproduction than would a large sit-and-wait species (Figure 4b). The same outcome should be observed if a widely-foraging species is larger and more efficient than a sit-and-wait species (Figure 4c). Furthermore, if a widely-foraging lizard is more efficient at foraging, it might forage for less time and reproduce more frequently than would a sit-and-wait lizard. Therefore, widely-foraging lizards could not only produce larger or more offspring in a single reproductive event, but they could also increase the number of such events throughout life.

**Figure 4.**
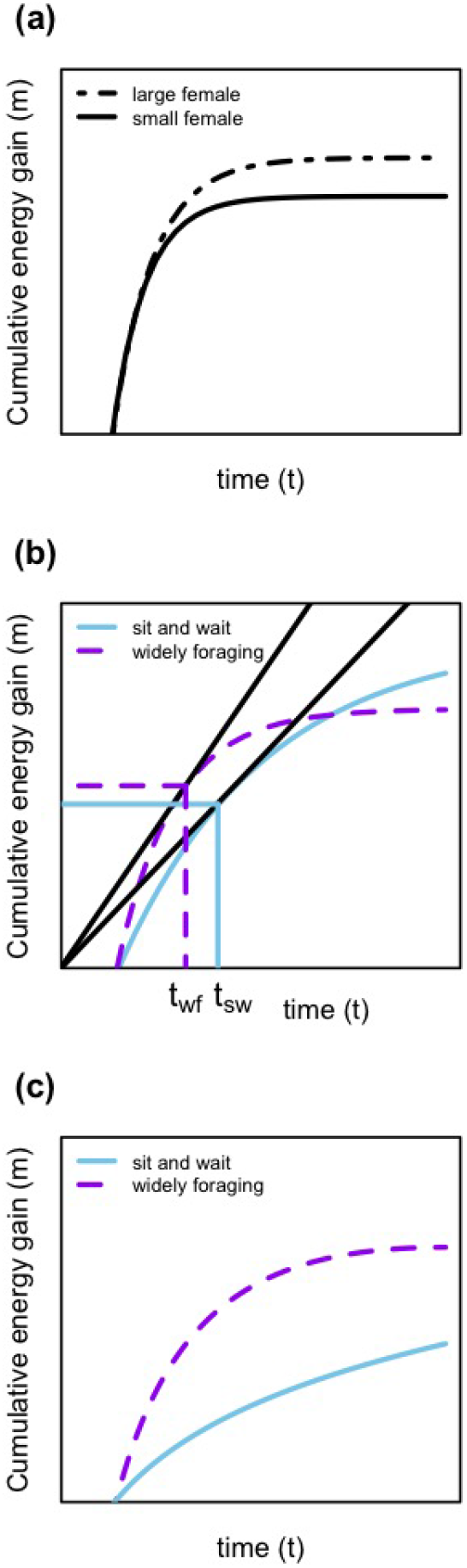
A theoretical model relating the cumulative energy gain for reproduction, *m*, as a function of time spent foraging, *t*, and maternal size. (a) Larger females reach their maximum capacity at a higher value of m than small females. (b) Widely-foraging females that are smaller but more efficient foragers may produce a greater reproductive effort than larger sit-and-wait females. (c) Widely-foraging females may also produce a greater reproductive effort than sit-and-wait females if they are both larger and more efficient foragers. *t_wf_* and *t_sw_* in (b) represent the optimal foraging time of widely-foraging females and sit-and-wait females, respectively.

Because foraging usually increases the risk of predation, individuals that require less time to accumulate resources should incur a lower chance of death. Schoener hypothesized that the energy accumulated from foraging increases monotonically toward an asymptote [53]. In such cases, foraging for twice as long would not result in twice the energetic return (Figure 4b). However, the rate of energy gain likely depends on foraging mode, as well as the abundance and distribution of prey. Consistent with this idea, widely-foraging lizards grew to a larger body size while foraging for less time than sit-and-wait foragers did [11]. The ability of widely-foraging lizards to forage more efficiently should reduce their risk of predation. Rapid growth by widely-foraging lizards might also cause them to outgrow the gape limitations of predators [54, 55]. Therefore, if body size is critical to outperform predators and outcompete conspecifics, the optimal pattern may be to grow to the maximal size that leads to the greatest reproductive effort.

The foraging-mode paradigm is focused on dichotomous variation, yet plastic variation in foraging mode [56] may drive the evolution of the reproductive traits in some species. Recent studies have revealed that the foraging mode of an organism depends on the ecological context, such as the presence of predators, the abundance of prey, or the surrounding habitat [57, 58, 59, 60]. Such plasticity of behavior could precede rapid evolutionary change and local adaptation of the life history [61]. A mixed foraging strategy was represented by species in the superfamilies Anguimorpha, Gekkota, Iguania, and Scincoidea (Figure 2). Because organisms adopting a mixed foraging strategy are possibly exposed to a wide range of environments with different selective pressures, these species might actively select habitats that maximize their reproductive effort, indirectly resulting in local adaptation [61]. Our analysis, which provides some evidence that a mixed foraging strategy likely leads to the greatest reproductive effort, should encourage others to address the questions raised by our observations. Additionally, future analyses might consider temporal patterns of foraging associated with seasonal environments. For example, widely-foraging lizards in the Kalahari Desert consume more food during the summer and stop eating during winter, as they hibernate. By contrast, other species of lizards ambush prey during both seasons [62]. Evidence of this nature is still rare, revealing the need for long-term studies of the energetics of foraging modes within and among species.

Expanding our analysis to include additional life-history variables could resolve or even alter the observed patterns. For instance, in areas with long growing seasons, production of multiple clutches offsets the constraint of optimal egg size on clutch size, leading to a greater lifetime reproductive effort than a single clutch of either a few large eggs or many small eggs [63]. Similarly, age of first reproduction is a critical component of the life history. In risky environments, for example, a species of lizard should evolve a reduced age of first reproduction and greater reproductive effort [22]. Similarly, direct comparisons of the mortality rates of widely-foraging species versus sit-and-wait species are required to know whether foraging mode affects a lizard’s vulnerability to predation. Although this evidence would be difficult to obtain for hundreds of species, detailed studies of a strategic sample of species would complement the broad comparative analysis that we conducted. Ultimately, a combination of comparative and experimental analyses will be needed to develop a general perspective of how foraging mode shapes life-history evolution.

Our study presents the first evidence that the early shift in foraging mode—from sit-and-wait foraging to widely foraging—in the evolutionary history of lacertilians was likely accompanied by the evolution of a greater reproductive effort, specifically in large-bodied species of lizards (Figure 3). Yet, much variation in reproductive effort exits among species with the same foraging mode, such that ecologists must consider other factors that influence reproduction. Importantly, our study captured the effects of foraging plasticity on the reproductive effort of lizards. For instance, lizards that adopt a mixed-foraging strategy produced the greatest reproductive effort. Hopefully, our findings encourages others to investigate how foraging efficiency and predation risk associated with foraging modes influence the evolution of reproductive traits.

## Acknowledgements

We thank Mark O’Shea for providing us with lizard photos for data visualization. We also thank Gustavo Burin for his valuable comments on an early draft of our manuscript, and Camila Tabares for helping us edit some of our figures.

## Data accessibility

A fully reproducible workflow of the data analyses associated with our study, including R code and additional supporting material, is available in the following Dryad Digital Repository [64]: https://doi.org/10.5061/dryad.xd2547djc.

## Author Contributions

D.J.P. conceived the study and analyzed data. D.J.P, D.F.D, and M.J.A. drafted the manuscript. All authors revised the manuscript, approved the final version, and agreed to account for its content.

